# A charge-dependent phase transition determines interphase chromatin organization

**DOI:** 10.1101/541086

**Authors:** Hilmar Strickfaden, Ajit K. Sharma, Michael J. Hendzel

**Author notes:** Address for correspondence: Dr. Michael J Hendzel, Department of Oncology, University of Alberta, Cross Cancer Institute, 11560 University Avenue NW, Edmonton, Ab, Canada, T5M 2T6, Dr. Hilmar Strickfaden, Department of Oncology, University of Alberta, Cross Cancer Institute, 11560 University Avenue NW, Edmonton, Ab, Canada, T5M 2T6.

## Abstract

An emerging principle of cellular compartmentalization is that liquid unmixing results in formation of compartments by phase separation. We used electron spectroscopic Imaging (ESI), a transmission electron microscopy technology, to distinguish chromatin and nucleoplasmic phases of mammalian cell lines and their responses towards different environmental changes. We tested the hypothesis that charge-dependent phase separation mediated by the histone N-termini could explain the organization of chromatin. 3D images of nuclear chromatin with electron spectroscopic imaging (ESI) demonstrates that the amount of chromatin proximal to the interchromatin compartment (IC) differs between cell types, reflecting major differences in chromatin organization. These differences were lost when cells were treated overnight with a histone deacetylase inhibitor. We show that drastic, reversible changes in chromatin mixing or unmixing with the nucleoplasm/interchromatin space can be induced by modulating osmolarity of the medium or acetylation status of the chromatin. In vitro phase separation experiments demonstrated that chromatin separated from solution through a phase transition towards a more solid chromatin state.

## Introduction

Two key historical developments underpin our current understanding of the cell nucleus: the development of the light microscope and the genome-wide application of molecular biochemical techniques that map biochemical relationships to underlying DNA sequence and higher order nuclear structures. In the early days of light microscopy, the development of staining techniques allowed the identification of structures such as the nucleolus (Hertwig, 1876) and nuclear bodies, such as the Cajal body (reviewed in (Ogg and Lamond, 2002)) and Barr body (Barr and Bertram, 1949). Chromatin, on the other hand, was largely not visible outside of the mitotic phase of the cell cycle. With the introduction of transmission electron microscopy (TEM) (Knoll and Ruska, 1932) and its superior resolution, it became possible to directly visualize the molecular assemblies that comprise the cell. Although this has led to a good understanding of the structure of the cytoplasm including membrane systems, cytoskeleton, and composition of organelles, progress in understanding nuclear architecture was delayed by its highly compact and crowed nature.

An important advance in this understanding became possible by the development of methods for specifically contrasting RNA on ultrathin sections in the TEM, EDTA regressive staining (Bernhard, 1968). The application of this method led to the first appreciation of the division of the interphase nucleus into a chromatin and an interchromatin compartment by revealing that the interchromatin compartment is dominated by nucleoli and interchromatin granule clusters (IGCs); the latter correspond to the sites of poly(A) RNA and splicing factor accumulation rather than decondensed chromatin. Ribonucleoproteins (RNPs) and RNA containing “coiled bodies” or Cajal bodies could also be found in the nucleoplasmic phase of the nucleus between domains comprised of chromatin, while RNA-rich perichromatin granules and fibrils were shown to flank the chromatin (Fakan and Bernhard, 1971; Fakan and Bernhard, 1973; Monneron and Bernhard, 1969; Petrov and Bernhard, 1971). Pulse chase experiments with radioactive tritium labeled nucleotides revealed that RNA synthesis takes place in the perichromatin region (PR) - the interface between chromatin domains and interchromatin space(Vecchio et al., 2008). The RNPs of the transcripts are subsequently released into the interchromatin space where they are then eventually transported through the nuclear pores into the cytoplasm (Fakan and Bernhard, 1973). Studies on DNA replication (Fakan and Hancock, 1974; Fakan et al., 1972; Niedojadlo et al., 2011), DNA transcription (Vecchio et al., 2008) and DNA repair (Solimando et al., 2009) demonstrated that these processes are also happening in the PR. However, there are examples of actively transcribed genes that invade the interchromatin space. The Balbiani ring genes occur as highly decondensed puffs that encode highly transcribed proteins in polytene chromosomes in direct contact with the surrounding nucleoplasm. Additionally, genes have been inferred to loop into the interchromatin space in human nuclei based on their separation from the territory associated with their chromosome.

Recently, molecular methods that attempt to map 3D contacts and neighborhood arrangements within the genome such as Hi-C (Lieberman-Aiden et al., 2009) and GAM (Beagrie et al., 2017) in combination with methods that map the interaction between proteins and DNA (ChIP-Seq (Johnson et al., 2007) DamID (van Steensel and Henikoff, 2000)) and SPRITE (Quinodoz et al., 2018), have provided interesting insights into the topological organization of the eukaryotic genome. According to these findings, the genome is organized into loops forming topologically associated domains (TADs) with a remarkable evolutionary conservation (Dixon et al., 2012; Vietri Rudan and Hadjur, 2015) despite functionally important intra-TAD interactions, whose relationship with specific gene expression patterns of a given cell is still not well understood (Dixon et al., 2012). The loops are built by extrusion of the DNA with the help of architectural proteins, such as cohesin and condensin (Barrington et al., 2017; Vian et al., 2018) and then bridged at the stem by CTCF. Using these genome-wide molecular techniques, it was possible to independently demonstrate the existence of chromosome territories (Lieberman-Aiden et al., 2009) previously described by FISH (Cremer and Cremer, 2010). It also provided evidence for the existence of a transcriptionally more active A compartment that seems to assume more peripheral positions at the surface of CTs and a transcriptionally less active B-compartment, which seems to be preferentially surrounded by chromatin belonging to the A-compartment (Lieberman-Aiden et al., 2009). Fundamentally, both techniques report on spatial relationships that are captured following crosslinking with aldehydes and should provide consistent but complementary information. Models describing these results need to capture both the molecular contacts mapped by chromosome conformation capture methods and the structure that can be directly observed by light and electron microscopies.

A current model, called the ANC-INC model (Cremer et al., 2015), partitions chromatin into an inactive heterochromatic compartment, the INC, and an active, transcriptionally active compartment, the ANC. The ANC is built up from a dynamic system of interchromatin compartment channels lined by the perichromatin region (PR). IC-channels extend from the nuclear pores, permeate the peripheral layer of heterochromatin located beneath the lamina and expand into IC-lacunae with splicing speckles and other nuclear bodies. The lining of the PR interacts directly with the interchromatin compartment (IC). It is enriched in transcriptionally competent chromatin and serves as the major site of transcription. The largely transcriptionally inactive chromatin of the INC is more compacted and located more remote form the IC. Such an organization has the benefit that it enforces a direct interaction of the active chromatin (especially the PR) with transcription factors. Transcription factors are believed to be enriched within the IC and the formation of a PR may minimize the scanning process in order to promote regulatory fidelity in gene transcription.

Our objective was to test predictions of the ANC/INC model with respect to the chromatin-interchromatin space organization using electron spectroscopic imaging (ESI) (Aronova and Leapman, 2013; Bazett-Jones and Hendzel, 1999; Hendzel and Bazett-Jones, 1996; Strickfaden et al., 2015). ESI provides analytical, spatial information about atoms, such as phosphorus strongly enriched in DNA, and nitrogen, strongly enriched in proteins. This information has helped to deconvolute the interphase nucleus with respect to the spatial arrangements of DNA particularly enriched in chromatin domains and proteins enriched in the IC. We find that the nuclear topography differs significantly between cell types. The fractions of space occupied by the ANC and INC compartment differ slightly between cell lines but conform to the general principles of the ANC/INC model. We further find that nuclear organization is extremely sensitive to environmental and biochemical changes. Changes in osmotic stress (Albiez et al., 2006; Amat et al., 2018) or histone acetylation status (Taddei et al., 2001) can dramatically alter the organization of the chromatin and interchromatin compartments. Moreover, we also find that the isolated chromatin fibers within the nucleosplasm were heterogeneous across the range of cell lines and exist within a size range of 5 to 50 nm thickness rather than as discrete 10 or 30 nm fibers, consistent with recently published measurements using ChromEMT (Ou et al., 2017). In vitro experiments revealed that a phase transition of chromatin to a more solid state provides a mechanism to explain the organization of interphase chromatin and its dependence on histone acetylation status.

## Results

### Organization of the interphase nucleus across cell types

Figure 1 shows ESI images of a mouse embryonic fibroblast cell nucleus where semi-quantitative maps of phosphorus (red) and nitrogen (green) are employed to identify nuclear structures and compartments based on their composition. The figure shows more compact domains of chromatin around the nucleoli and the nuclear envelope. In mouse nuclei, particularly large areas of clustered heterochromatin can be observed as chromocenters – areas where the pericentric and telomere-associated heterochromatin of the telocentric mouse chromosomes aggregate. Less compact chromatin can be found at the interface of the chromatin domains to the interchromatin compartment (IC) – a protein dominated environment, that extends from the nuclear pore complexes to the interior of the nucleus and occupies the space in-between the chromatin domain. This IC contains fibrillar nucleoplasmic protein structures, nuclear bodies, mRNPs and interchromatin granule clusters (nuclear speckles) (see Introduction).

**Figure 1:**
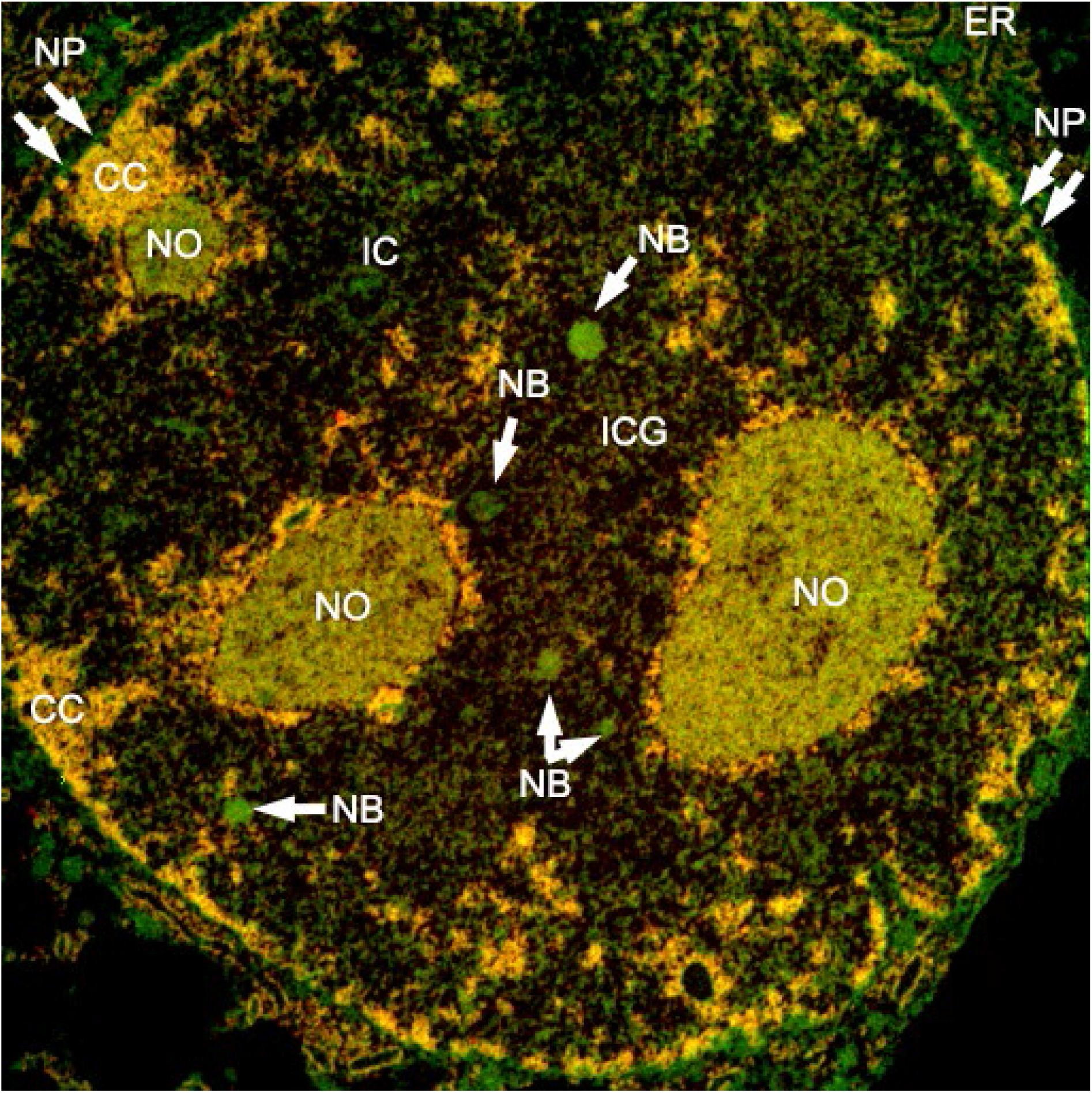
ESI of a mouse embryonic fibroblast nucleus. Elemental maps of phosphorous and nitrogen in the sample are ideal to show the distribution of nucleic acids (phosphorous is displayed in red) and protein (nitrogen is displayed in green). The image shows the nuclear envelope with nuclear pore complexes (NP) the lamina nucleoli (NO), nuclear bodies (NB) chromocenters (CC) chromatin (shown in yellow color) and the interchromatin space (IC).

Employing this type of electron microscopy, we assessed the relationship between the chromatin and interchromatin space to examine how the organization of the interphase nucleus differs across cell types that strongly differ with respect to their functional states (Figure 2 and SFig 1). ESI images reveal strong differences in the nuclear topography of nucleic acids (false colored in red) and proteins (false colored in green). Nuclei from cultured mouse embryonic fibroblasts (C3H/10T1/2), human neuroblastoma cells (SK-N-SH), and chimeric human/mouse hybridoma cells carry dense chromatin networks that can have chromatin structures with diameters greater than one hundred nanometers. These are co-aligned with broad interchromatin channels and lacunae, which harbor interchromatin granule clusters and nuclear bodies. In contrast, the nuclear organization of HeLa cells is characterized by much more dispersed “open” chromatin where the individual chromatin fibers are well mixed with the interchromatin compartment and it is hard to find chromatin structures with diameters greater than that of single fibers. We quantified the organization of the interphase nucleus by defining each pixel as INC, ANC, or IC based on distance from the surface of the chromatin region. This evaluation revealed marked differences in the distribution of chromatin between INC and ANC compartments that are consistent with the visible differences between the cell lines (Figure 2).

**Figure 2:**
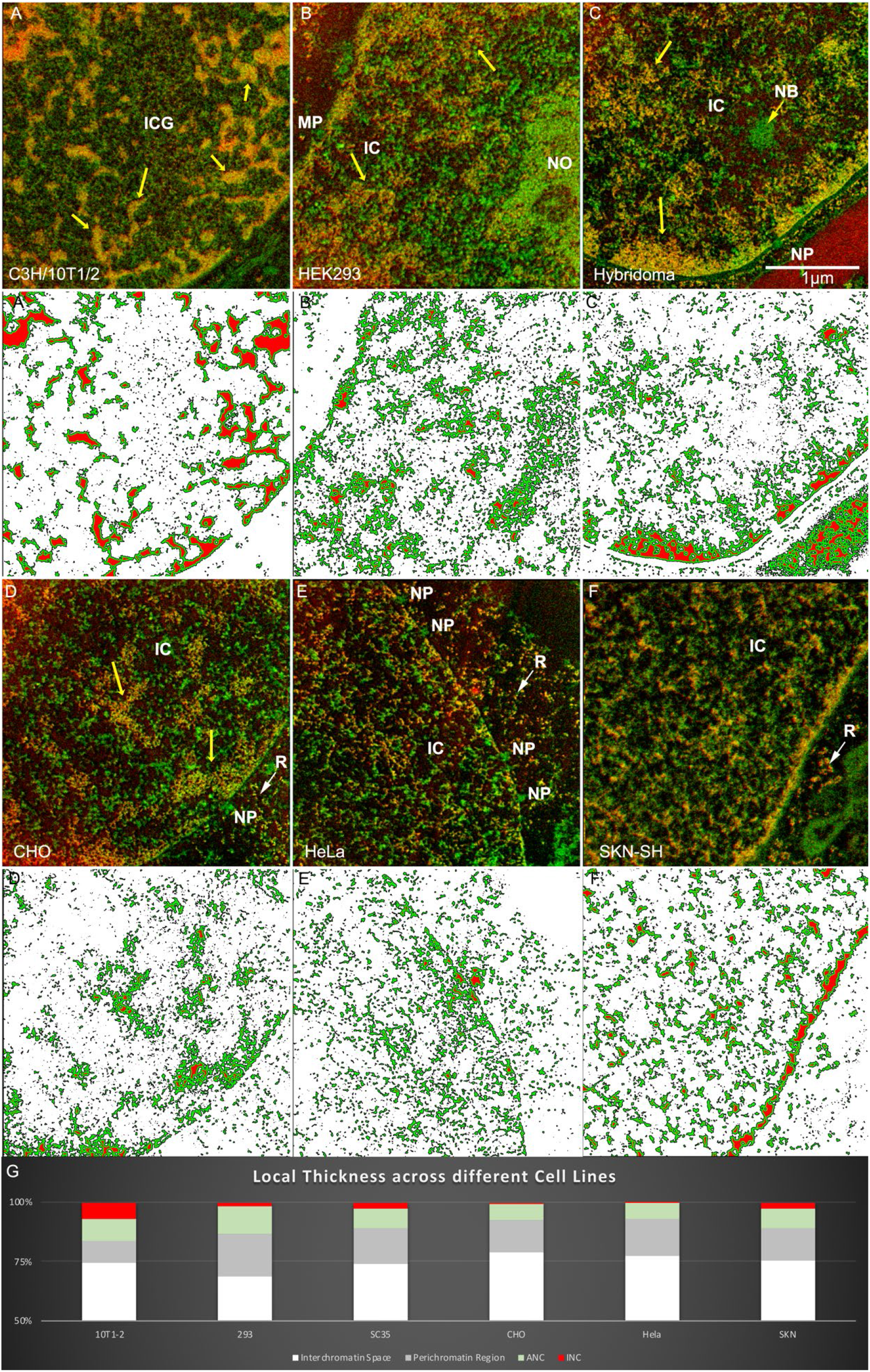
Individual cell lines show significantly different nuclear configurations of the chromatin. (A-F) Net phosphorus and net nitrogen images were collected and merged images shown for each cell line. Below the ESI pictures (A’-F’), the local thickness of the structures composed by nucleic acids (phosphorus) are measured. While white shows the regions free from nucleic acid (Interchromatin Space), grey shows the chromatin in immediate contact (5nm) with the interchromatin space (Perichromatin Region). The areas painted in green are areas that are not further away from the interchromatin space than 20nm (Active Nuclear Compartment (ANC)). Red shows areas that are more than 20 nm from the interface of the interchromatin space and the chromatin phase (Inactive Nuclear Compartment (INC)). G) Bar charts that show the proportions between interchromatin space (white), chromatin, directly in contact to the interchromatin space (grey), proximal (green) and distal chromatin (red).

### The effect of hypertonic and hypotonic treatment on nuclear organization

It has been established by fluorescence microscopy that chromatin organization can be dramatically and reversibly altered at time scales in the order of a minute by changing the osmolarity of the tissue culture medium (Albiez et al., 2006) SFigure 2 and SMov1. In order to study the reorganization of the chromatin and the interchromatin space, we incubated cells in culture medium with different osmolarities and fixed cells of mouse embryonal fibroblasts (C3H/10T1/2), the human neuroblastoma cell-line SK-N-SH and U2OS cells (osteosarcoma). Figure 3 and SFig 3 (middle column) shows ESI images of nuclei of the same three cell lines that were cultured and fixed under isotonic (control) conditions (290 mOsm). Figure 3 and SFig 3 (left column) shows ESI images of the nuclei from cells that were incubated and fixed in hypertonic medium (570 mOsm). Within one minute, this treatment prompted massive changes of the nuclear architecture. The chromatin organization that arises is consistent with the expectation that increased osmolarity induces an enhanced phase separation of the chromatin from the interchromatin compartment and nucleoli. In line with this expectation, the nucleoli seen after hypertonic treatment were rounder and often detached from chromatin by a surrounding interchromatin space, whereas nuclei prepared under isotonic conditions carried irregularly shaped nucleoli with numerous dense, directly attached chromatin structures. In addition, peripheral heterochromatin was detached from the lamina. Note how well defined the lamina is in these images. Under normal preparation conditions, it is hard to resolve the lamina from the lamina associated chromatin. Following hypertonic treatment, the IC shows a marked increase in the abundance of nuclear bodies compared to nuclei from cells fixed under isotonic conditions. These nuclear bodies exclude chromatin and RNAs but the proteins that comprise these bodies are unknown. This result suggests that phase separation events are also taking place within the nucleoplasmic protein pool. When cells were fixed after incubation in hypotonic medium (~140 mOsm), see Figure 3 and SFig 3 (right column), a more subtle reduction of higher order chromatin configurations were noted. The chromatin appeared more dispersed and pervaded by finely dispersed interchromatin channels.

**Figure 3:**
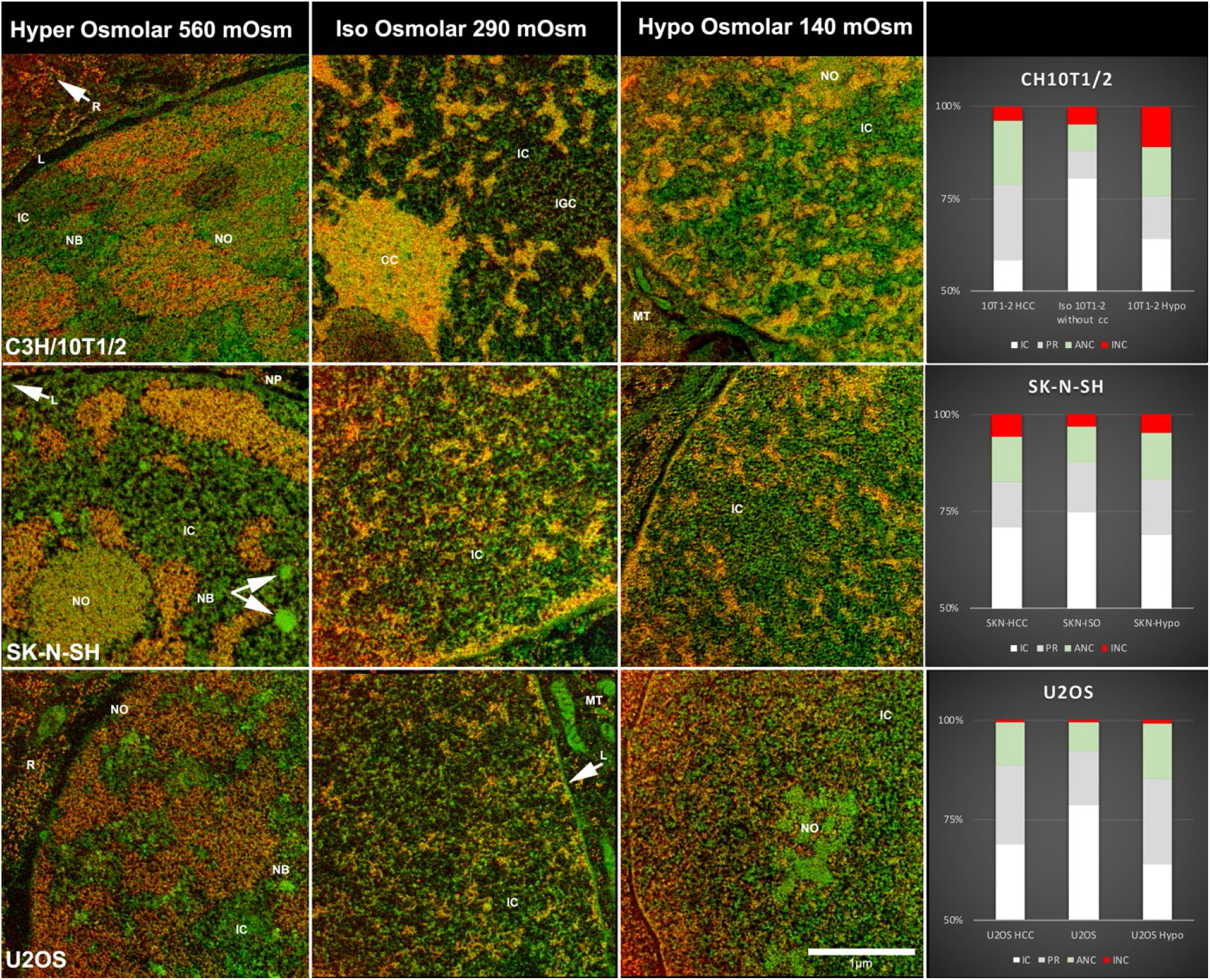
Nuclear organization is very sensitive to changes in osmolarity. ESI images of ultra-thin (50 nm) sections C3H/10T1/2 (upper row), SK-N-SH (middle row) and U2OS (lower row) cells were incubated and fixed under different osmotic conditions (Hyper osmolarity 560 Osm (left column), Iso osmolarity 290 Osm (middle column) and Hypo osmolarity 140 Osm (right column)). Net phosphorus and net nitrogen maps were collected and superimposed. The bar graphs on the right show the relative amounts of interchromatin space (white), chromatin at the interface to the interchromatin space (grey), proximal (green) and distal (red) chromatin.

To obtain a quantitative assessment of these differences, we measured the fractions of IC, perichromatin layer, ANC and INC. We found an increase in the in the amount of chromatin and a decrease of the area occupied by the IC in cells treated with hyper-and hypo-osmolarity compared to cells fixed under iso-osmolar conditions.

### Effect of histone hyperacetylation on nuclear organization

Osmolarity is a mechanism of modulating the structure of the chromatin by changing the concentration of ions in the nucleoplasm. Cells can implement a similar phenomenon in a physiological environment by altering the charge on the chromatin fiber itself. One way to achieve this is through the neutralization of positively charged lysine residues of histone proteins through reversible posttranslational acetylation. To test the influence of charge on the organization of interphase chromatin, we treated cells with the histone deacetylase inhibitor trichostatin A (TSA) for approximately 18 hours (Figure 4). At high magnification (Figure 4E-F), we observe the interchromatin space to be comprised of proteinaceous fibrils (green) with variable diameters (ranging from about 9 to 18 nm), together with a chromatin fiber network (red) (ranging from 8 to 24nm). The chromatin is dispersed, similar to, although more dramatically and less structured than, what is observed after hypoosmotic treatment. This is consistent with previous studies that measured smaller “pore sizes” and “increased chromatin accessibility” using fluorescence correlation spectroscopy and nuclear microinjection of differently sized, fluorescently labeled dextrans in HeLa cells (Gorisch et al., 2005). Figure S4A-F shows striking similarities with ESI images recorded from other undifferentiated cells such as pluripotent stem cells (supporting online Figure 4), HeLa cells and U2OS cells. The separation of the IC and the chromatin compartment is less distinct in these cell types, suggesting that the large majority of chromatin is well mixed with the interchromatin compartment.

**Figure 4:**
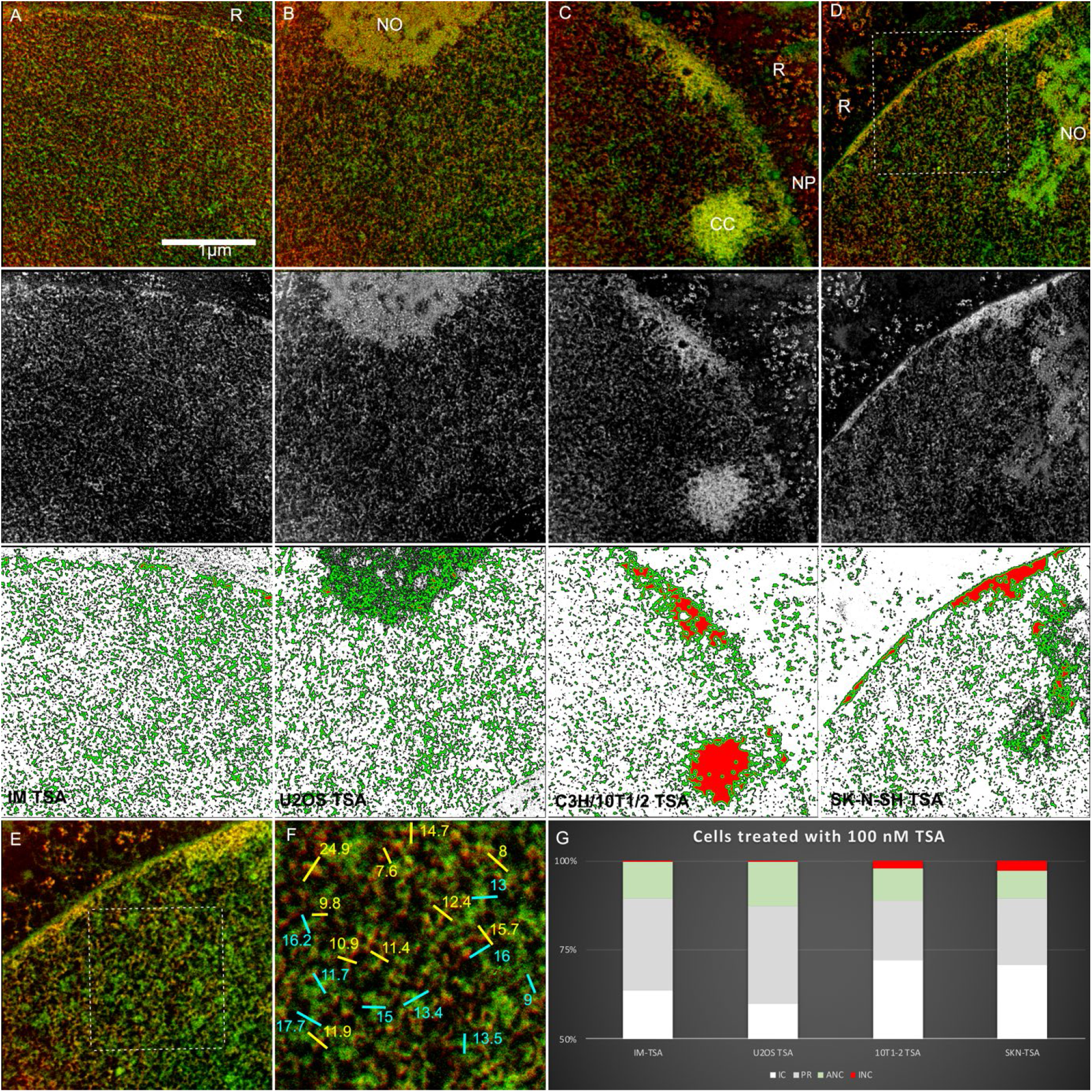
Histone acetylation dramatically alters chromatin organization. **A-D:** ESI images (upper) phosphorous elemental maps (middle), thickness analyzed maps (lower) of 50 nm ultrathin sections show nuclei of (A) Indian Muntjac, (B) U2OS, (C) C3H/10T1/2 and SK-N-SH cells (D) treated with 100nM TSA for 24h. **E-F:** Decondensed chromatin fibers and interchromatin fibrils shown in D at higher magnification and with line scans in F. **G:** Bar charts showing the amounts of Interchromatin Space (white) – chromatin at the interface to the interchromatin space (grey) and proximal (green) and distal (red) chromatin. For the measurements the areas that were occupied by nucleoli, chromocenters were excluded from the evaluation.

Mouse chromocenters are assemblies of pericentric heterochromatin from telocentric mouse chromosomes that assume an approximately spherical organization in interphase mouse nuclei. Minimizing surfaces is typical for phase separated membrane-less compartments in the nucleus that form by liquid-liquid phase separation (Erdel and Rippe, 2018). Untreated mouse cells (C3H/10T1/2) show the typical IC/chromatin arrangement (see Figure 1 and 2A). Upon TSA treatment both chromocenters and the peripheral layer of heterochromatin underneath the nuclear lamina retained a more compact organization that excludes interchromatin space (Figure 4A-D). This observation suggests that a phase-separated liquid or a more solid structure persists even after prolonged incubation with TSA. TSA treatment blurred the otherwise visible distinction between chromatin and interchromatin space. A quantitative comparison of the fractions of interchromatin space and the different classes of chromatin revealed a clear shift towards chromatin that is in direct contact with the ANC. In our study of TSA-treated mouse cells, most chromatin classified as INC was confined to peripheral chromatin or chromocenters.

In order to further assess the relationship between histone acetylation and the chromatin structure that we observed across cell lines under normal growth conditions, we analyzed the acetylation state of the histones using acetic acid-urea-Triton X-100 polyacrylamide gel electrophoresis. This enables the resolution of acetylated species of histones based on the neutralization of the positive charge on the target lysine residue. In vitro experiments have revealed that as acetylation increases, resistance to forming assemblages that sediment upon centrifugation increases (Tse et al., 1998). These experiments revealed that it was the total amount of acetylation rather than the acetylation of specific acetylation sites that dictates the solubility of chromatin in vitro. Consequently, we were interested in determining the extent of multiply acetylated histone species. We selected U2OS as an example of a nucleus with very dispersed chromatin and SK-N-SH and C3H/10T1/2 cells as examples of cells with a more “condensed” phenotype. The U2OS cells, as expected, had the highest amount of multiply acetylated species (Figure 5). This is evident in both the H3 and the H4 regions of the gel. The H4 region of the gel is shown separately using Sypro Ruby protein stain, which was more sensitive than Coomassie Blue. Interestingly, the C3H/10T1/2 cells had a state of acetylation that was intermediate between the U2OS and SK-N-SH cells. However, a second feature was strikingly different that provides an explanation for the apparently enhanced phase separation of the interphase chromatin. C3H/10T1/2 cells showed a markedly greater histone H1 content. Since histone acetylation and histone H1 have been shown to act in concert to promote chromatin insolubility *in vitro,* this result is consistent with the observed morphology and implicates both histone H1 content and histone acetylation in regulating the state of the interphase chromatin outside of the heterochromatin domains.

**Figure 5:**
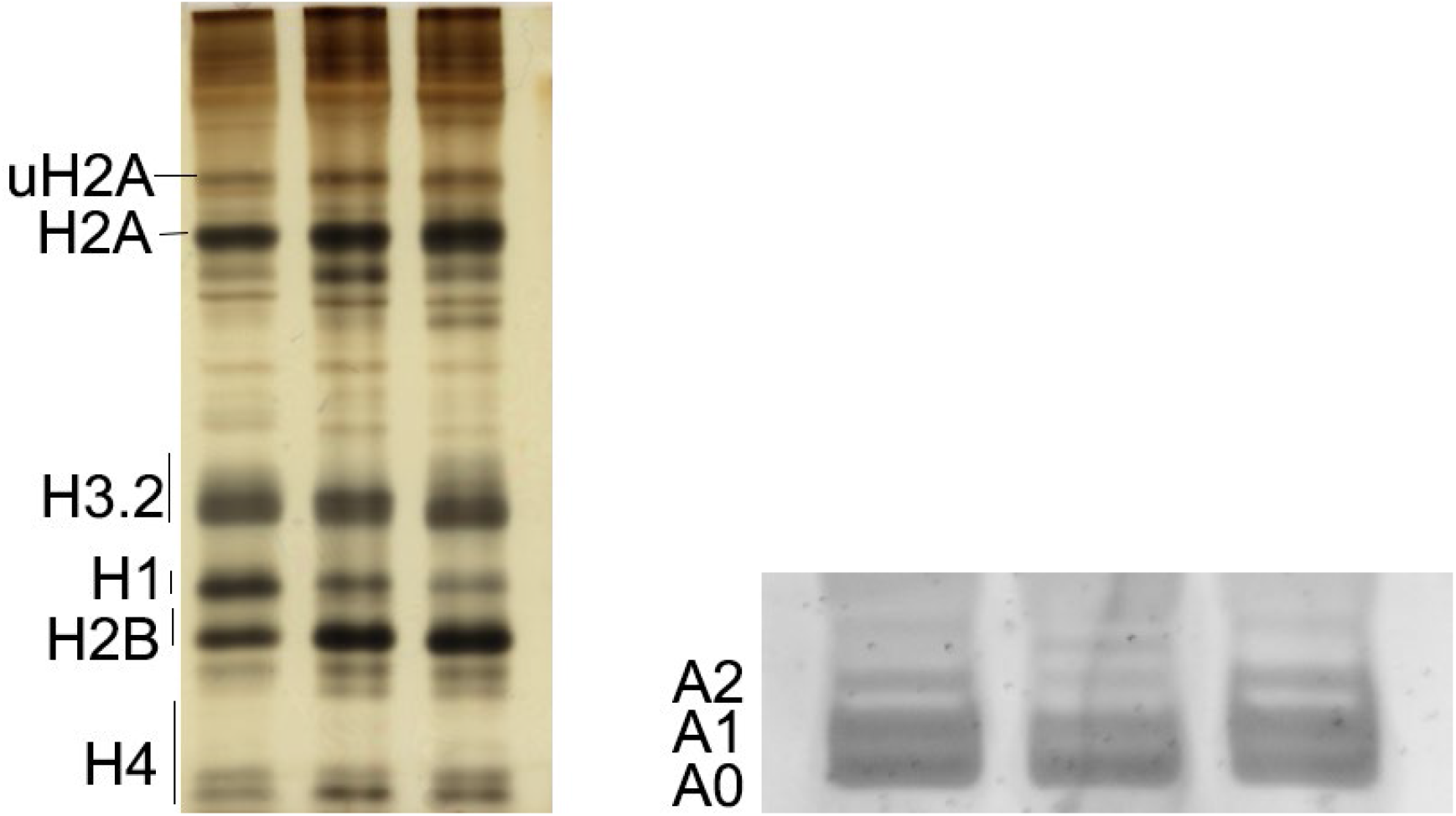
Histone composition and acetylation state of chromatin. Acid-soluble proteins were resolved by acetic acid-urea-Triton X100 15% polyacrylamide gel and stained with silver (left) and Sypro Ruby (right). Histones H2A, H2B, H3.2, H4 and H1 are labelled. The lower panel shows the corresponding H4 region where individual acetylated species are indicated.

### Chromatin fiber structure across cell types, acetylation status, and osmotic treatment

The nature of the structural organization of chromatin beyond the nucleosome particle has been an ongoing source of controversy. In *in vitro* experiments, 10nm ‘beads on a string’ 10 nm chromatin fibers are observed in the absence of H1 and at low ionic strengths. In the presence of low mM concentrations of divalent cations or histone H1, 30 nm thick chromatin fibers are observed (reviewed in (Hansen, 2002)). With the exception of sea urchin sperm and chicken erythrocytes (Woodcock, 1994), the existence of the 30 nm fiber as an in vivo structure, however, has been challenged (Fussner et al., 2011; Maeshima et al., 2010; Ou et al., 2017). A study of the condensed chromatin of chromocenters in mouse cells with ESI-tomography revealed only extended fibers of approximately 10 nm and no evidence for a higher-order fiber (Fussner et al., 2012). EM Tomography of DRAQ-5 stained and photooxidized chromatin (ChromEMT) also did not provide evidence for a 30 nm fiber. Rather, only chromatin fiber diameters between 8 nm and 24 nm were observed (Ou et al., 2017). Consequently, we wished to assess the chromatin fiber diameter amongst these cell lines with markedly different chromatin organizations.

We have performed fiber measurements on our phosphorous images of nuclei from different species under control isotonic conditions, following TSA treatment or following hyper or hypotonic treatment (Figure 6A). In order to minimize measurement errors, we have identified cytoplasmatic ribosomes in every sample in order to illustrate the range of diameters that an approximately 25 nm diameter structure presents as in thin sections. Consistent with previous reports, we were not able to identify 30 nm thick fibers regardless of the cell system or the environmental conditions used at the time of fixation. The fact that we could not find evidence for a common well-ordered folding of the chromatin beyond the nucleosome level is fully consistent with previously reported results obtained with the ChromEMT approach (Ou et al., 2017). While we noted larger regions of chromatin where multiple fibers appear to contribute to the observed chromatin structure, these are comprised of separate nucleosome chains rather than a single chain adopting a hierarchically folded state. These observations are consistent with the model that was presented in (Ricci et al., 2015) where amorphous aggregates of chromatinized DNA fibers described as “clutches” arise from inter nucleosome interactions (SFig 5).

**Figure 6:**
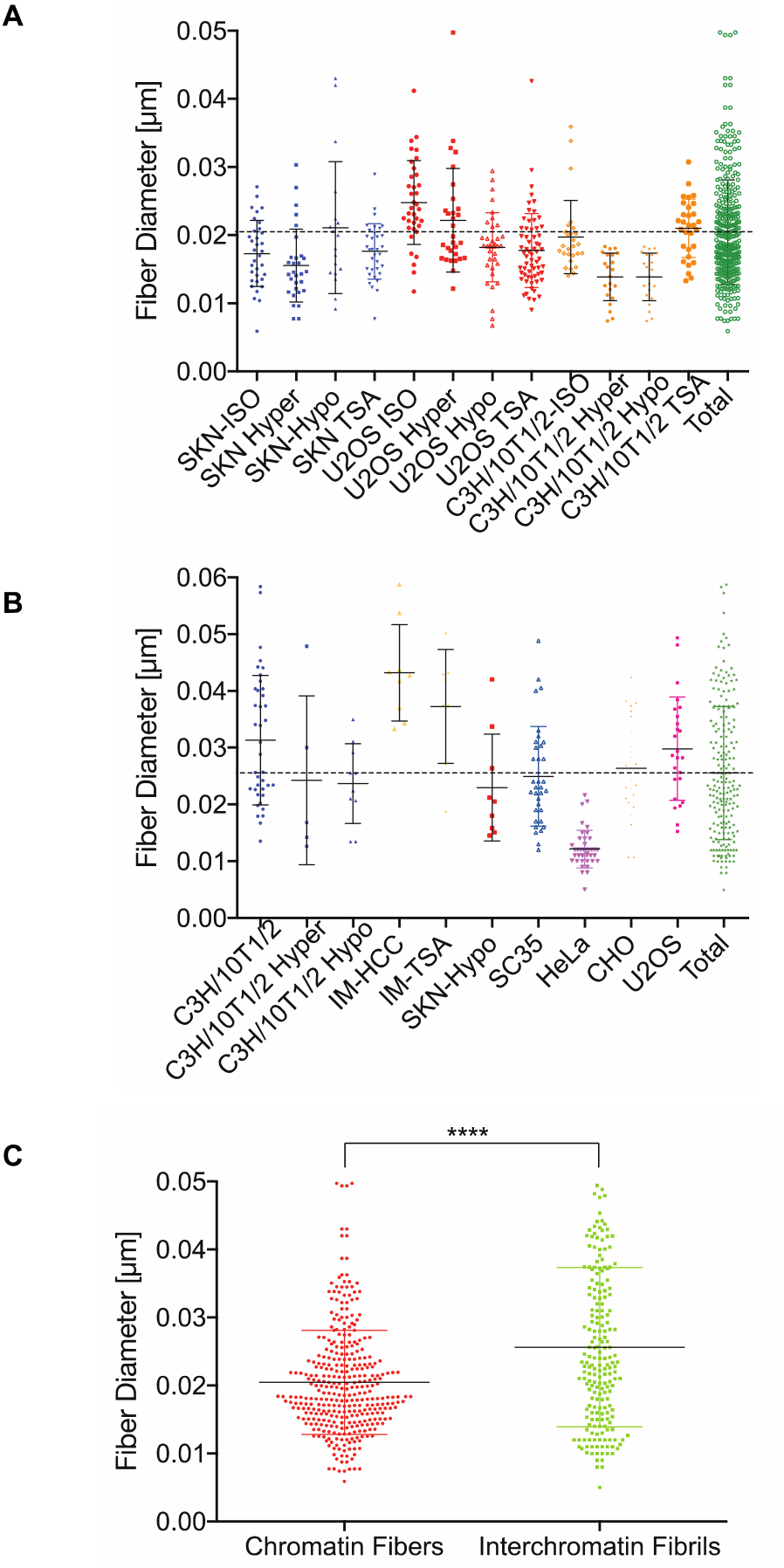
The physical dimensions of chromatin and protein filaments of the interphase nucleus. A: Measurement of the chromatin fiber diameters in different cell lines. B: Measurement of the interchromatin protein fibrils measured in the interchromatin space. C: The diameters of the chromatin fibers vs. interchromatin fibrils are statistically different in size.

### Chromatin undergoes a charge-dependent phase transition in vitro

It is well established that the majority of cellular chromatin is insoluble in buffers containing 2 mM or higher magnesium concentrations and/or monovalent ions at concentrations between 100 and 200 mM. However, the reorganization of chromatin to enable its collection by centrifugation has not been examined directly. Consequently, we performed chromatin fractionation (Figure 7a) and examined the organization of chromatin in a buffer containing 100 mM NaCl and MgCl_2_ (Figure 7c). Figure 7b shows a phase diagram revealing that as the magnesium concentration increases, there is a rapid drop in the fraction of the chromatin that remains in solution between 0.5 and 2 mM and then a gradual decrease in soluble chromatin at higher magnesium concentrations. This is consistent with the chromatin being sensitive to physiological concentrations of magnesium. Figure 7c shows the organization of this unmixed chromatin visualized by the addition of Hoechst 33258 and fluorescence microscopy. Importantly, the chromatin formed networks based on the association of smaller domains with each other rather than forming ever larger phase-separated liquid domains. The irregular morphology of the structure in the liquid environment is consistent with a solid, rather than a liquid, nature to the unmixed chromatin and suggests that phase transition, rather than phase separation, is what accounts for the ability to fractionate interphase chromatin.

**Figure 7:**
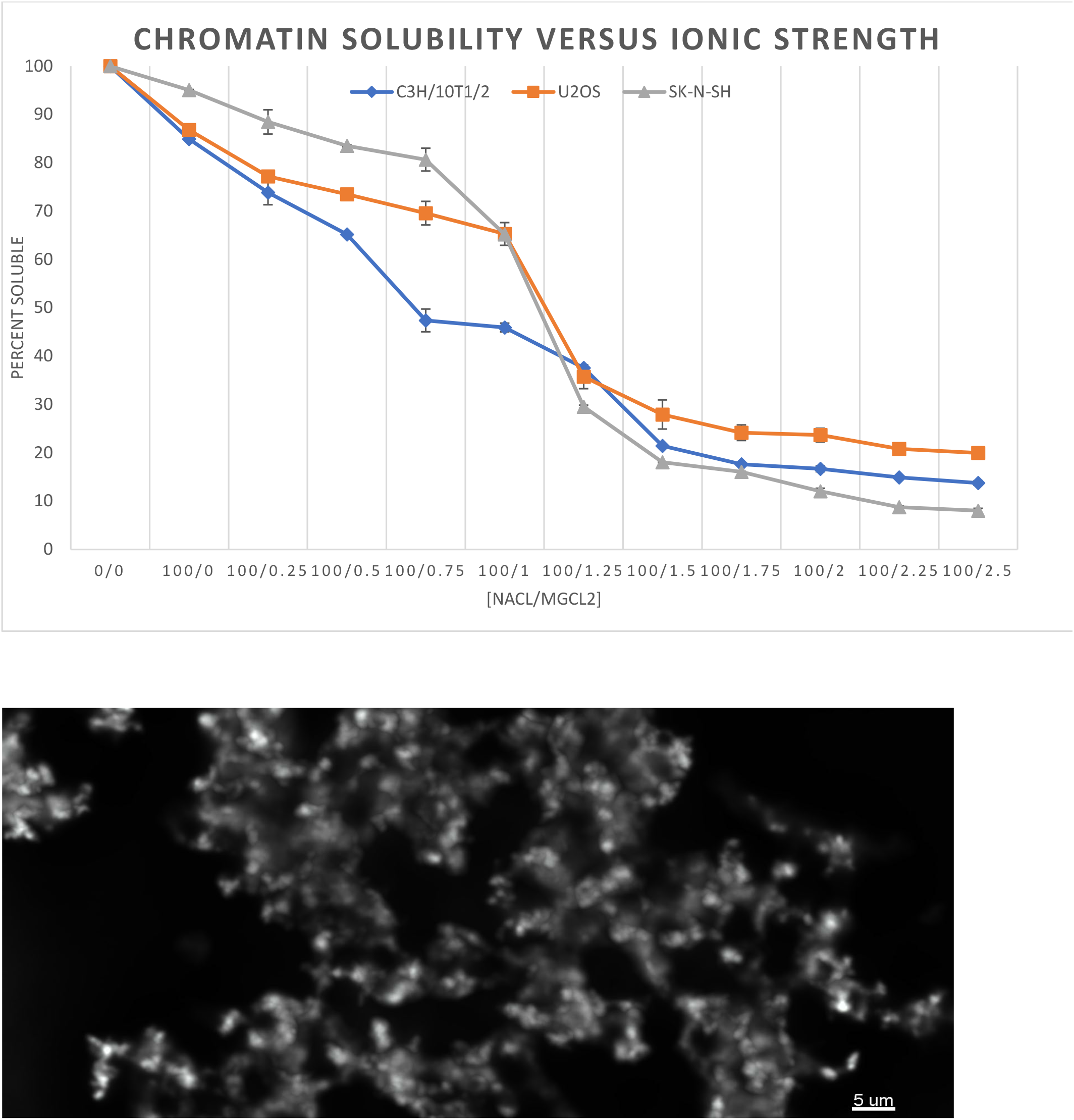
Chromatin undergoes a phase transition in the presence of physiological concentrations of magnesium ions. Upper: Phase diagram showing the chromatin solubility of the cell lines C3H/10T1/2 (blue), U2OS (red) and SKN-SH (green) as a function of increasing MgCl2 concentration. Lower: Chromatin in 5 mM MgCl_2_ was mixed with 1 μM Hoechst and visualized by fluorescence deconvolution microscopy. Images were acquired every 200 nm and the deconvolved image stack was then cropped to eliminate out of focus information near the top and bottom of the image series. The total depth of the image shown is 8 μm.

### Unmixed chromatin domains are poorly accessible to nucleoplasmic proteins

To test whether size exclusion of nuclear proteins is responsible for the different accessibilities to chromatin domains, we transfected C3H10T1/2 cells with fluorescent nuclear proteins with a range between 14 kDa (miniSOG) and 293 kDa (CBP-GFP) (see Figure 8 and Table S1). Since EGFP (27 kDa) and mCherry (28.8 kDa) have a similar molecular weight to DNase I (31 kDa), we transiently expressed these proteins in MEFs and examined their distribution in living cells using Hoechst as a DNA specific stain to define the distribution of the chromatin (Figure 8). The nucleolus, which has been characterized as a phase-separated compartment (Feric et al., 2016), served as a reference. EGFP and mCherry are clearly depleted in the nucleolus (~0.1 relative fluorescence intensity). The EGFP and mCherry, however, were also not uniformly distributed in the nucleoplasm outside of nucleoli, but clearly depleted in the chromocenters (~0.05-0.1 relative fluorescence intensity). We also tested nuclear proteins of increasing molecular weight to determine if their soluble nucleoplasmic pools could access chromocenters. Rad52-GFP (73 kDa), AP2b-GFP (77.5), HDAC-1 GFP (82 kDa) and CBP-GFP (292 kDa) showed similar distributions within the nucleoplasm (Figure 8). FCS analysis revealed the existence of significant amounts of monomeric populations of the GFP-tagged expression products (e.g. 58% for HDAC1-GFP). Hence, the ability of these proteins to incorporate into complexes is not defining the reduced accessibility. Interestingly the 14 kDa fluorescent protein miniSOG, which is about half the molecular weight of DNase 1, was also excluded from chromocenters and nucleoli and showed similar nucleoplasmic distributions in the interchromatin compartment.

**Figure 8:**
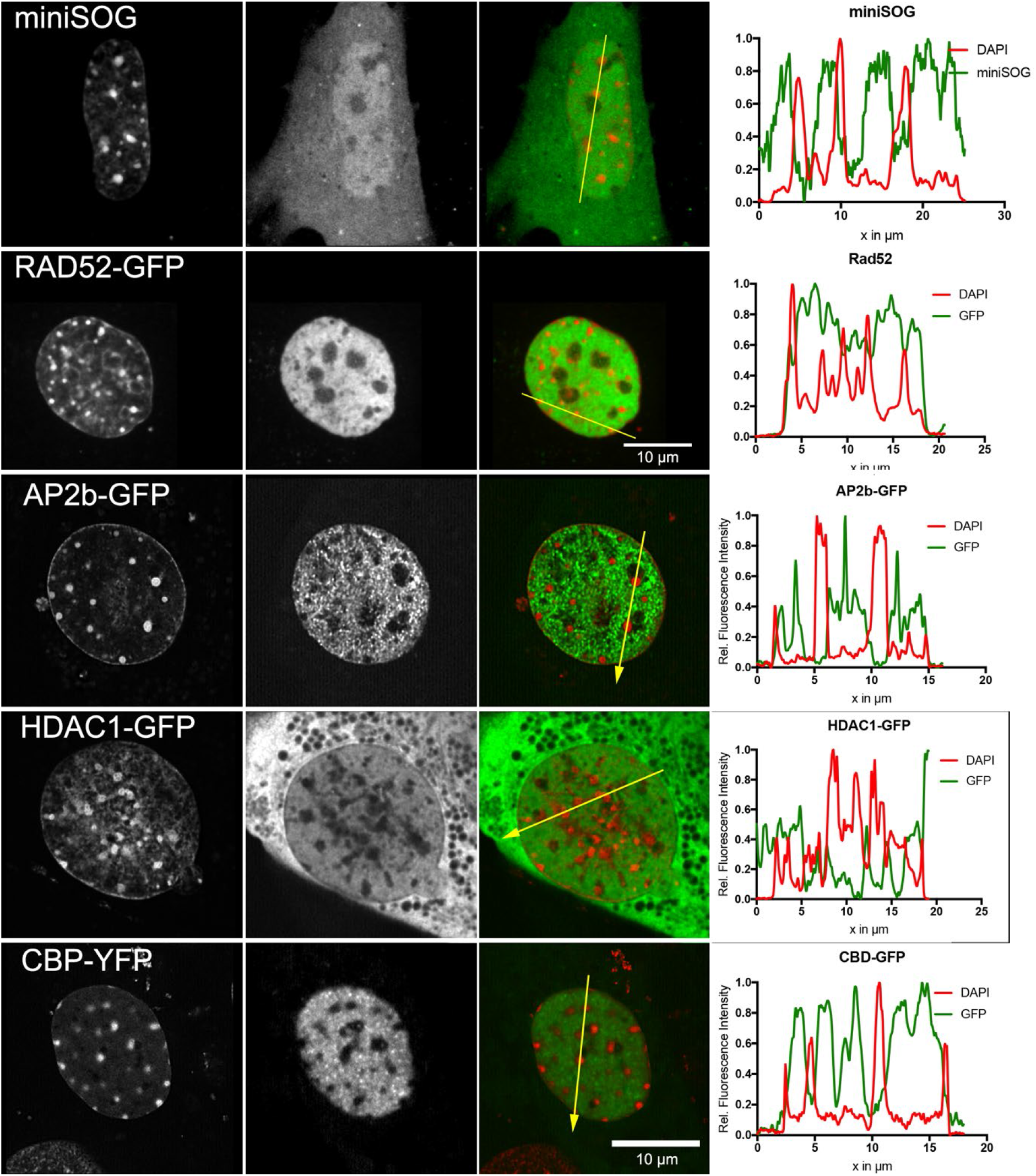
Accessibility of chromatin to proteins of increasing molecular weight. C3H/10T1/2 cells transfected with the fluorescent proteins indicated in the left column. Leftmost column is Hoechst staining, the center images show the tagged protein alone while the image series on the right shows the composite images. A line scan showing the intensity profiles along the yellow line indicated in the third column is shown on the right.

Due to the lack of the z-resolution (~700 nm), it was not possible to demonstrate a clear depletion of these fluorescent proteins in chromatin dispersed throughout the ANC as compared to chromatin present at much higher compaction within the INC. However, we clearly noted fluctuations in intensity of fluorescent proteins that correlated with increases and decreases in chromatin density suggesting that higher chromatin compaction limits the accessibility of otherwise freely diffusing nucleoplasmic proteins (Figure 8). We sought to verify this with a small native nuclear protein, a splicing factor, SC35, fused to EGFP, and PML fused to dsRed (Figure 9). PML bodies are dense protein-rich domains that are also believed to be phase separated (Mitrea and Kriwacki, 2016). PML bodies also show depletion of SC35-GFP. It can be seen that SC35-GFP is similarly excluded from chromocenters. It is notable that chromocenters appear significantly less dense/more porous by ESI than nucleoli but exclude these nucleoplasmic proteins similarly. This argues against simple physical occlusion as the mechanism of excluding small nucleoplasmic proteins.

**Figure 9:**
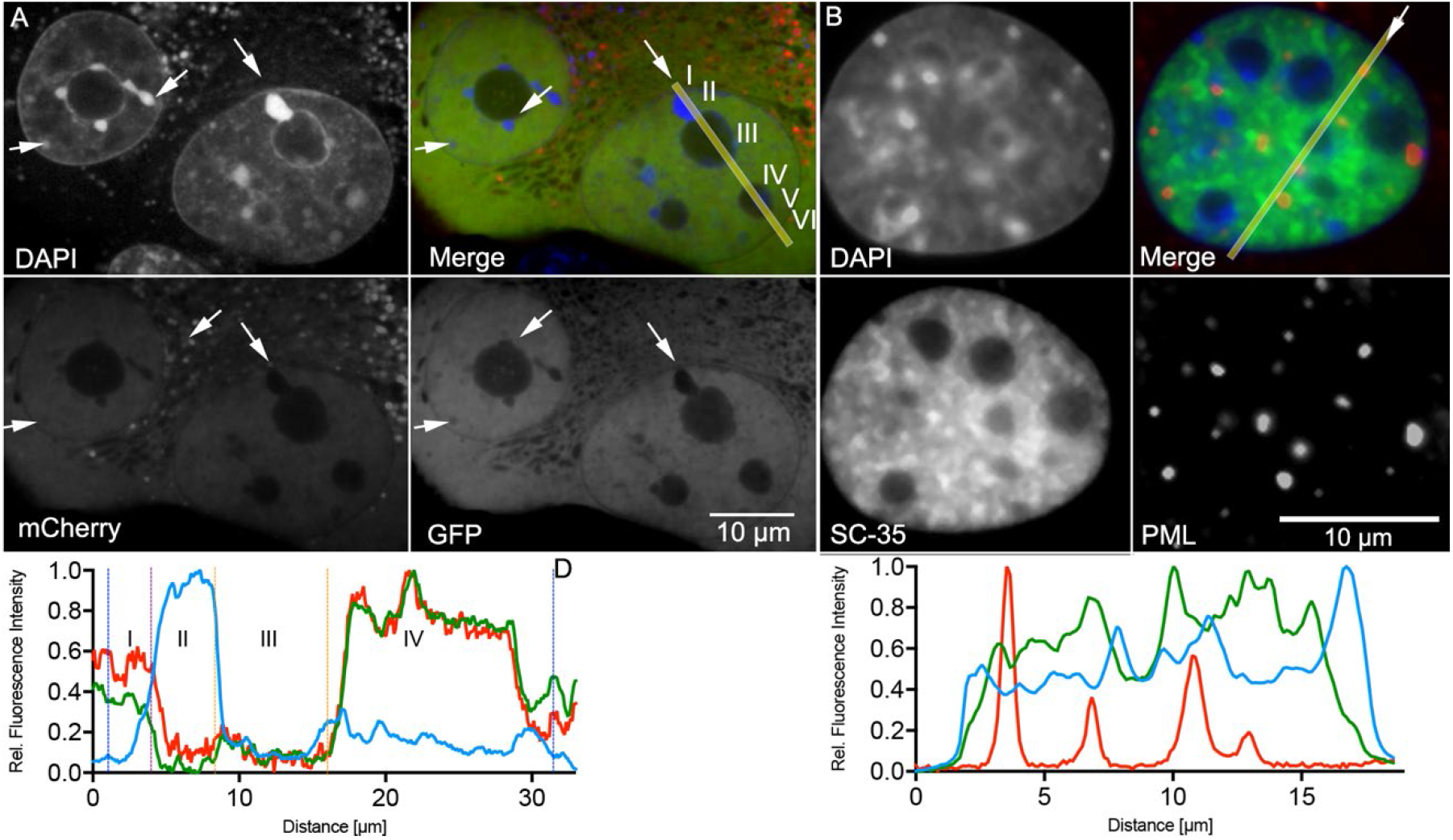
Accessibility of small molecules in the interchromatin space is limited by chromatin and nuclear bodies. **A:** C3H/10T1/2 nucleus transfected with SC35-GFP and PML-dsRed and stained with DAPI shows exclusion of the nucleoplasmic SC35 from areas that are occupied by chromatin (especially chromocenters) nucleoli and (to lesser extent by) PML bodies. Line-Scan shows the fluorescence intensity profiles of DAPI, SC-35-GFP and PML-dsRed. B: C3H/10T1/2 nuclei transfected with eGFP and mCherry and stained with DAPI shows that fluorescence intensity of the small fluorescent proteins is reduced at sites of dense chromatin (especially chromocenters (arrows)) and nucleoli. Line-Scan shows the fluorescence intensity profiles of DAPI, eGFP and mCherry.

## Discussion

The organization of interphase chromatin and its relationship to the accessibility of the genome has been a longstanding and unresolved question (Branco and Pombo, 2006; Cremer et al., 2015; Maeshima et al., 2010). In this work, we employed electron spectroscopic imaging (ESI) to image nucleic acids and protein in unstained, ultrathin sections at subnucleosomal resolution. Because of its ability to resolve different classes of biomolecules and facilitate the identification of isolated chromatin fibers in a manner that clearly distinguishes them from any filamentous protein structures, ESI is particularly well suited to study the interface between chromatin and the interchromatin space. In addition to profound, cell line-specific morphological differences and the interchromatin space forming an intricate channel network flanked by chromatin, we found characteristics in nuclear organization that are consistent with chromatin mixing and unmixing with the nucleoplasm/IC being an important regulatory mechanism to control access to DNA.

We have also shown that environmental changes of osmolarity or drugs affecting the acetylation state of the histones has a profound effect on the mixing/demixing of chromatin with the interchromatin space/nucleoplasm. In light of this, we argue that changing ionic or osmotic conditions (Albiez et al., 2006; Visvanathan et al., 2013) affects chromatin organization by modulating mixing/demixing of chromatin. Under hypertonic conditions, the chromatin in all observed cell types forms large assemblies of dense chromatin. Under these conditions, the chromatin in mouse nuclei became condensed to an extent that the chromocenters were no longer distinguishable from the surrounding chromatin. This treatment generated large interchromatin space lacunae with an increased number of nuclear bodies and detachment of the peripheral heterochromatin from the nuclear lamina. We interpret this as unmixing from the nucleoplasm to form chromatin-rich domains.

Recently, phase separation was proposed to be an organizing principle of pericentric heterochromatin, which was shown to form a subcompartment in the nucleus that differs in protein composition and biophysical properties from the surrounding interphase chromatin (Strom et al., 2017). This stands in contrast to models that explain accessibility of chromatin by size exclusion caused by differently sized pores that result from variations in the density of folding (Nozaki et al., 2017). Interestingly we found that pericentric heterochromatin and the peripheral heterochromatin remained compact and retained its characteristic structure after addition of TSA to mouse embryonal fibroblasts for an extended period of time, extending what was previously shown by fluorescence microscopy (Taddei et al., 2001). This is in stark contrast to the remainder of the chromatin, which was driven to a fully decondensed state following TSA treatment. The maintenance of these heterochromatin structures following TSA treatment may be related to observations made with molecular methods like DamID (van Steensel and Belmont, 2017) and Sat4C (Wijchers et al., 2015), which were able to identify heterochromatin compartments such as PADs (pericentromere-associated domains) (Wijchers et al., 2015) and LADs (van Steensel and Belmont, 2017) as distinct from each other and the rest of the genome based on the relative frequency of interactions within the domain versus outside of the domain.

Using small fluorescent proteins, both native and exogenous, we further demonstrated that demixing of chromatin associated with mouse chromocenters results in a high depletion of nucleoplasmic proteins. Since we used very small proteins that are soluble in the nucleoplasm, this accessibility is expected to be at least as restricted for the much larger regulatory proteins and complexes involved in transcriptional activation. In light of current evidence that extended chromatin fibers exist even within constitutive heterochromatin and taking into account the failure to demonstrate the presence of a transcriptionally repressive chromatin structure composed of folded, well-defined 30 nm thick fibers, we propose that acetylation-driven chromatin mixing and deacetylation driven chromatin demixing act as a basic mechanism that defines the differences in accessibility of active and inactive gene sequences. The finding that active transcriptionally regulating elements like promoters and enhancers and other highly transcriptionally active regions like the Balbiani Rings are regions of highly acetylated histones (Turner et al., 1990) plus the fact that histone acetyltransferases are basic components or closely associated with the Pol II machinery (Struhl, 1998) supports this view. The large diversity in chromatin configurations that we have shown between different cell-types, acetylation states, or osmolarity are examples of features that have not been adequately captured by molecular techniques. The ANC-INC model is postulating that the surfaces of CDCs contain the perichromatin region of the genome that is active and mixing with the interchromatin compartment, whereas chromatin within the interior of CDCs has little to no access to the interchromatin channel network, helping preserve its inactive state.

If these phase separated chromatin domains poorly mix with the IC/nucleoplasm, this could provide a biophysical mechanism to establish a nuclear segregation into distinct, co-localized ANC and INC compartments (Albiez et al., 2006; Cremer et al., 2015).

Chromatin fibers are highly crosslinked through histone H1 and the core histone N-terminal tails. Accordingly, we propose a phase state change from a more liquid state of chromatin within the ANC to a more solid state within the INC rather than a phase separation of two liquid states by liquid-liquid or liquid-polymer phase separation. This is evident in the organization of the chromatin after it demixes from solution in the presence of magnesium. Even after prolonged incubation (48 hours), the chromatin assemblies remain physically stable, have irregular borders, and do not fuse into larger spherical domains, which would be indicative of a liquid state. Consequently, the results of the present study are in line with a large body of literature demonstrating that chromatin demixes from solution in physiological buffers, where it can then be collected by centrifugation. The extensive history of analysis of the influence of ionic strength on chromatin conformation has been reviewed in detail (Hansen, 2002). Classical biochemical fractionation studies revealed that most chromatin in insoluble in buffers containing mM concentrations of magnesium (Perry and Chalkley, 1982) and monovalent salt concentrations between 100 and 200 mM KCl or NaCl (Rocha et al., 1984). In purified systems, magnesium induces the oligomerization (phase transition) of reconstituted nucleosomal arrays but phase separation in the presence of monovalent ions requires histone H1. Similarly, the solubility of reconstituted chromatin fibers in 150 mM NaCl in the presence of increasing amounts of histone H1 is dependent upon histone acetylation (Ridsdale et al., 1990). It is this selective solubility of acetylated olignonucleosomes that is the historical basis for employing phase separation (salt insolubility) as a tool for isolating fractions enriched in active genes in classical chromatin fractionation procedures (Henikoff et al. 2009).

In chicken erythrocytes, highly expressed genes could be detected in salt soluble highly acetylated chromatin compartments (Jahan et al., 2016) Since these studies clearly illustrate that chromatin is at least poised to phase transition in ionic conditions similar to what is found in cells, it is not surprising that evolution ‘exploited’ this physical property to functionally organize the genome within the nucleus (Cremer et al., 2018). Acetylation has recently been shown to regulate the phase separation of DDX3X in the cytoplasm through a similar mechanism to what is described here, consistent with the hypothesis that charge neutralization of lysines in intrinsically disordered domains can regulate phase separation (Saito et al., 2019) Notably, the local regulation of magnesium concentration by hydrolysis of ATP was shown recently to participate in chromosome formation (Maeshima et al., 2018) indicating that regulating ionic strength is employed physiologically to regulate chromatin structure.

## Acknowledgements

We want to thank the Cross Cancer Institute Cell Imaging Facility for technical support and access to the transmission electron microscope and Bahaar Sehgal for helping with the sample preparation. This work was supported by operating grants from the Canadian Cancer Society Research Institute and the Canadian Institutes of Health Research.

## Author Contributions

The project was conceived by H.S. and M.J.H. H.S. designed the experiments, collected the image data and performed the quantification. M.J.H. contributed to experimental design.

Ajit K. Sharma performed the chromatin fractionation and in vitro solubility experiments. All authors contributed to writing the manuscript.

## Declaration of Interests

The authors declare no competing interests.

**Supplemental Information**

## Experimental Procedures

### Cell Culture

Cells were incubated at 37°C in a saturated water atmosphere at 5% in the respective growth media (see Table 1).

|  |  |  |
| --- | --- | --- |
| SK-N-SH | Neuroblastoma (Human) | DMEM 10% FBS |
| C3H/10T1/2 | Mouse Embryonal Fibroblast | Alpha MEM 10% FBS |
| U2OS | Osteosarcoma (Human) | DMEM 10% FBS |
| U2OS miniSOG mCherry | Osteosarcoma (Human) | DMEM 10% FBS |
| HeLa | Cervix carcinoma (Human) | DMEM 10% FBS |
| V79 | Hamster lung fibroblasts | DMEM 10% FBS |
| CHO | Chinese Hamster Ovary | F12 10% FBS |
| IM | Indian Muntjac | Ham's F-10 medium, 80% 20% FBS |
| HEK-293 | Human Embryonal Kidney | DMEM 10% FBS |
| anti-SC35 (ATCC CRL-2031) | Hybridoma | RPMI 10% FBS |

### Sample preparation for electron microscopy

Cells growing in 35mm MatTek dishes were fixed under iso-osmolar conditions in Sørensen buffer (290 mOsm) containing 2% glutaraldehyde for 30 min and then dehydrated in an ethanol series consisting of a 50%, 70%, 90% and a 98% step for 30 min with continuous agitation. After dehydration the cells were embedded into LR-White resin by first incubating them in a 1:1 mixture of LR-White and 98% ethanol for 12h and then incubating them in pure LR-White for another 12h. The dehydration and infiltration steps were done with continuous agitation and then the medium was replaced with fresh resin before curing them in an oven at 65°C for 24h as described in (Strickfaden et al., 2015).

The cured blocks were cut on a Leica ultramicrotome UC7. Floating 50nm thick sections were picked up on copper grids and an approximately 2 nm layer of carbon was deposited on the surface of the sections before inspecting and imaging them in the TEM.

### Hypo- and Hypercondensation Experiments

Hypo- and hypercondensation experiments were performed according to Albiez et al. (Albiez et al., 2006).

To fix cells under hypo-or hyperosmolar conditions, the cells were incubated in medium that had an osmolarity of 140 mOsm and 570 mOsm, respectively before fixing them in Sørensen containing 2% glutaraldehyde that had an osmolarity of 140 mOsm and 570 mOsm, respectively.

The fixed samples were further processed as described in the previous section.

### Hyperacetylation experiments

For the hyperacetylation experiments the cells were incubated for 24h in 100 nM TSA before they were fixed and prepared as described above. In-vivo chromatin accessibility analysis by expression of fluorescent nucleoplasmic proteins.

C3H10T1/2 cells were transfected the day before the observation with plasmid constructs coding for GFP-tagged proteins of different sizes that occur in the cytoplasm (see Table S1). 20 min before the observation the cells were stained with Hoechst 33342 (total concentration 10 μg/ml). Mid-sections of nuclei were imaged on a Perkin-Elmer Ultraview spinning Disk microscope attached to a Zeiss Axiovert 200M, a Zeiss Axiovert 200M widefield microscope and a Nikon A1R si HD laser scanning confocal microscope by recording the Hoechst and the GFP channel.

Line scans through nuclei were generated with ImageJ. The measured intensities were normalized and plotted with GraphPad Prism.

### Electron Spectroscopic Imaging

ESI was performed at a JEOL 2100F transmission electron microscope with a LaB6 filament operating at 200 kV. A Tridiem GIF post column spectrometer (Gatan) was used to the to record the elemental maps of phosphorus and nitrogen. Phosphorous and nitrogen maps were recorded at a magnification of 8500x. For the phosphorous map, electrons for the pre-edge images were collected at 120 eV with a slit width of 20 eV. The post-edge images were collected at 175 eV with a slit width of 30eV. For nitrogen maps pre-edge images were collected at 358 eV and post-edge images at 447 eV with a slit width of 35 eV. Pictures were processed as described in (Strickfaden et al., 2015)

### Fiber measurements

Fiber measurements were performed on phosphorous maps showing nuclear chromatin and on nitrogen maps showing interchromatin fibrils. Line scans were generated in ImageJ (Rueden et al., 2017) perpendicular to phosphorus- or nitrogen-rich fiber-like structures in the picture. The intensity profiles of the chromatin fibers and the interchromatin fibrils were measured in ImageJ using a self-written macro that determined the FWHM of the intensity profile based on a gaussian fit (see SFig 4). As an internal reference cytoplasmatic ribosomes were measured in order to compensate for focus-related changes in magnification. Their average diameter of a Ribosome was assumed to be 25 nm and the average of the ribosome measurements were normalized to this value.

### Measurements of the local thicknesses of nucleic acid structures in the nucleus

Phosphorous ratio maps of nuclei were smoothened in ImageJ by applying a median filter with a kernel of 3×3 pixel twice and then the picture was binarized using the Otsu method. An Euclidean distance map (EDM) was generated from the processed phosphorous map using the built-in binary processing tools of ImageJ. For quantitation and better visual inspection, the generated EDM was coloured with a look-up table that subdivides the pictures into the following classes: free-space – (white – no nucleic acid signal), Interface to the interchromatin space: (0-6 nm) (black – Perichromatin Region), chromatin behind the perichromatin layer but proximal to the interchromatin space 9-17nm from the surface (green ANC (Active Nuclear Compartment)) and distal nucleic acid area regions > 17nm (red – INC (Inactive Nuclear Compartment)). The histogram of the EDMs in areas inside the nuclei (except for chromocenters and nucleoli) were used to count the pixels that fell in each of the abovementioned categories and the proportions were calculated for each representative picture and displayed in a bar stacked column graph.

### Histone isolation and Acetic acid-Urea-Triton gel electrophoresis of histones

Murine fibroblast cell line (10T1/2) were cultured in α-MEM supplemented with 10% fetal bovine serum (FBS) at 37°C and 5% CO2 whereas U2OS and SKN-SH were cultured in Dulbecco’s modified Eagle’s medium (DMEM) with 10% FBS. Histones were extracted as described earlier (Hendzel et al., 1998) with few modifications. Harvested cells were washed twice with PBS, suspended in lysis buffer and incubated for 10 min at 4°C. The lysed cells were centrifuged at 5000 RPM for 10 min. Nuclei pellet were suspended in 0.2 M H_2_SO_4_ (6 vol.) followed by vigorous mixing at 4°C. After centrifugation at 16000 RPM at 4°C, supernatant was kept at −20°C overnight for precipitation of histones in acetone. The histone pellet obtained after centrifugation was washed with 50 mM HCl in acetone followed by washing with chilled acetone. Dried histone pellet was suspended in 0.1% β-mercaptoethanol in H2O and stored at −20°C. Total histone proteins were resolved on AUT-PAGE as described earlier by (Hendzel et al., 1998). Resolving gel contained 15% acrylamide, 0.1% Bis, 8 M urea, 1 M acetic acid, 50 mM NH4OH, 0.5% w/v Triton X-100, 0.5% TEMED and 0.0004% riboflavin 5’-phosphate. Stacking gel contained 4% acrylamide, 0.16% Bis, 8 M urea, 1 M acetic acid, 50 mM NH4OH, 0.5% TEMED and 0.0004% riboflavin 5’-phosphate. Total histones (10 μg) were loaded in each well and electrophoresis running buffer was 1 M acetic acid and 0.1 M glycine. The histone samples were electrophoresed at constant voltage of 150 V for 5 hr. After electrophoresis, the gels were stained with SYPRO Ruby protein gel stain (S12001, molecular probes Invitrogen).

### Preparation of Chromatin fragments and chromatin solubility assay against MgCl2

Nuclei were isolated by lysing cells in buffer pH 7.5 (10 mM Tris-Cl, pH 7.5, 10 mM NaCl, 3 mM MgCl_2_, 10mm sodium butyrate, 250 mm sucrose and 0.25% V/V of NP-40. Nuclei were washed two times with same lysis buffer and nuclei were collected by centrifugation at 5000 rpm for 10 minutes. Nuclei were resuspended at 50 A_260_ units per ml in digestion buffer containing 2 mM CaCl_2_ and were digested with 25 U/ml of micrococcal nuclease at 37°C for 20 min in digestion buffer (15 mM Tris−Cl pH 7.5, 15 mM NaCl, 250 mM sucrose, 2 mM CaCl_2_, 60 mM KCl, 15 mM β-mercaptoethanol, 0.5 mM spermidine, 0.15 mM spermine, 0.2 mM PMSF, protease and phosphatase inhibitors). The reaction was stopped by the addition of EGTA to 10 mM and nuclei were collected by centrifugation at 5000 rpm for 10 min. Nuclei were resuspended and lysed in 10 mM EDTA for 30 min on ice. The EDTA soluble chromatin was separated from insoluble nuclear material by centrifugation at 10000 rpm for 15 min. The isolated soluble chromatin was dialysed overnight against 1 mM Tris-Cl (pH 8.0) and 0.1 mM EDTA at 4°C. Soluble chromatin after dialysis recovered from dialysis bag and soluble dialysed chromatin was used to precipitate against various concentration of MgCl_2_ at 4°C. Chromatin solubility was determined by leaving a solution overnight at 4°C in buffer (1 mM Tris-Cl (pH 8.0) and 0.1 mM EDTA) at various concentration of MgCl_2_ and then centrifuge at 12500 rpm for 15 min with subsequent determination of the optical density of the supernatant at 260 nm.

### Fluorescence Correlation Spectroscopy

Measurements were performed on a Zeiss LSM 770 confocal microscope using an environmental chamber (37°C, humidified atmosphere containing 5% CO_2_) equipped with a ConfoCor 3 (APD) extension. Cells were grown in a MatTek 35 mm dish in phenol red free medium and measured the day after transfection with the respective expression construct coding for the GFP-tagged proteins shown in Table 1. Before the measurement the thickness of the coverslip at the respective position was measured by the total reflection of the laser beam and the 1.2 N Water immersion objective lens was adjusted accordingly. The ZEN-FCS plug-in was used to conduct the measurements, which were repeated 10 times per measurements over the period of 10 seconds. The curve fitting predicting a two-component model generated the best fits with the smallest residues.

---

**Supporting Online Figure 1:**
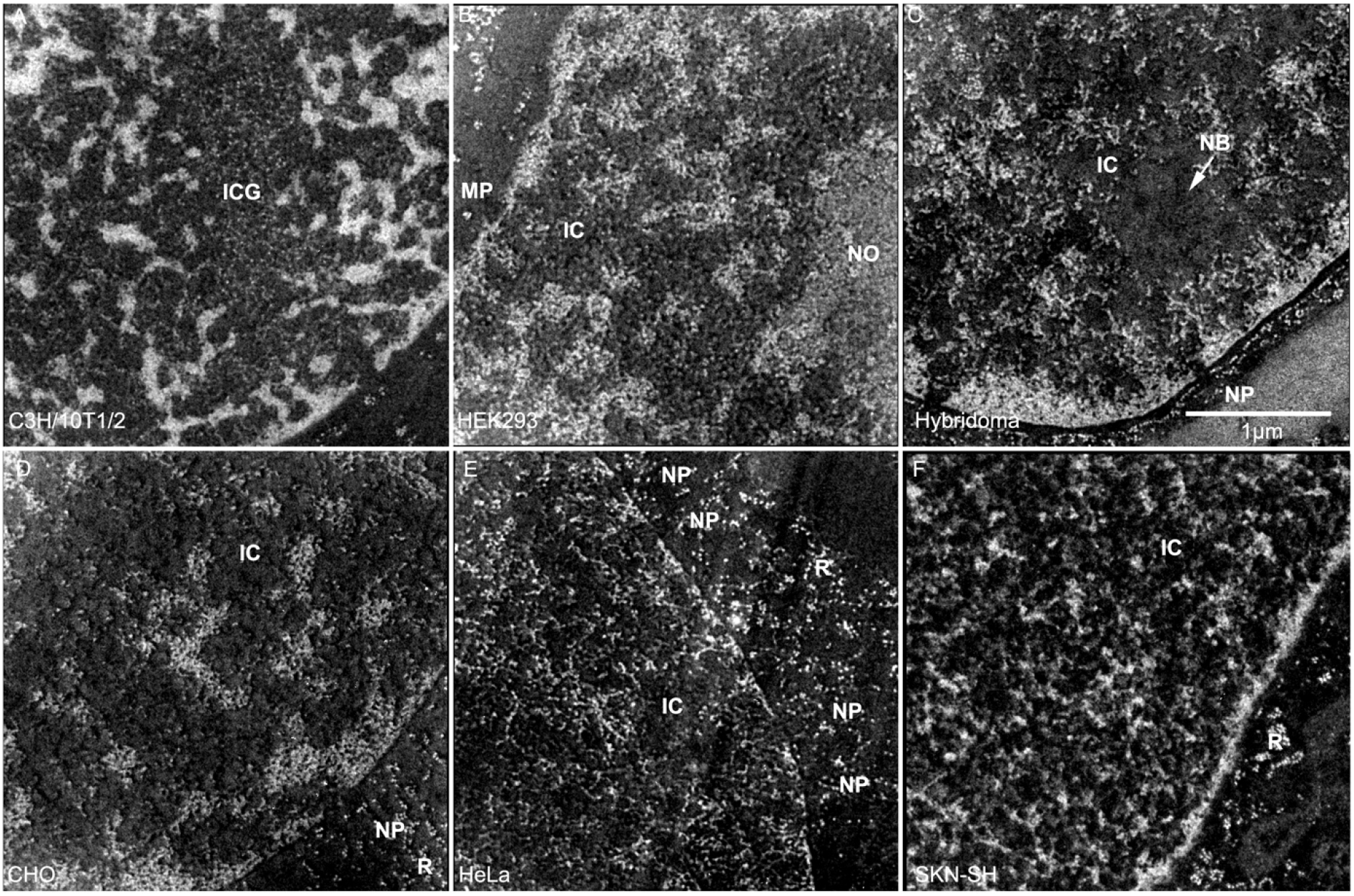
Phosphorous maps of the pictures shown in Figure 2

**Supporting Online Figure 2:**
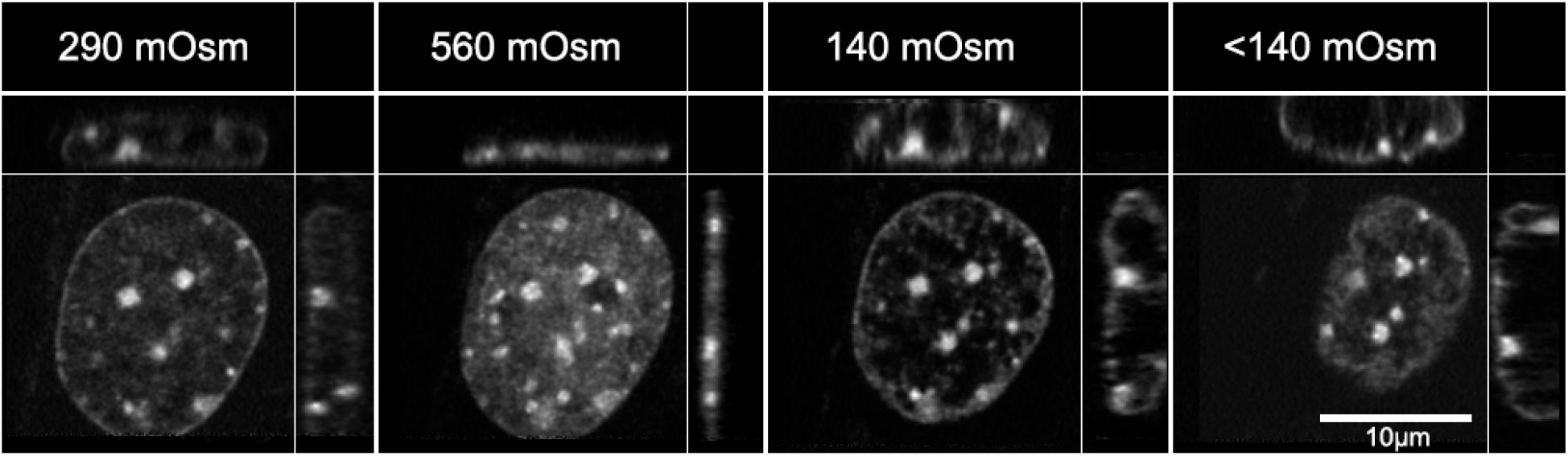
Reversible morphological changes in a C3H/10T1/2 nucleus induced by changes in osmolarity.

**Supporting Online Figure 3:**
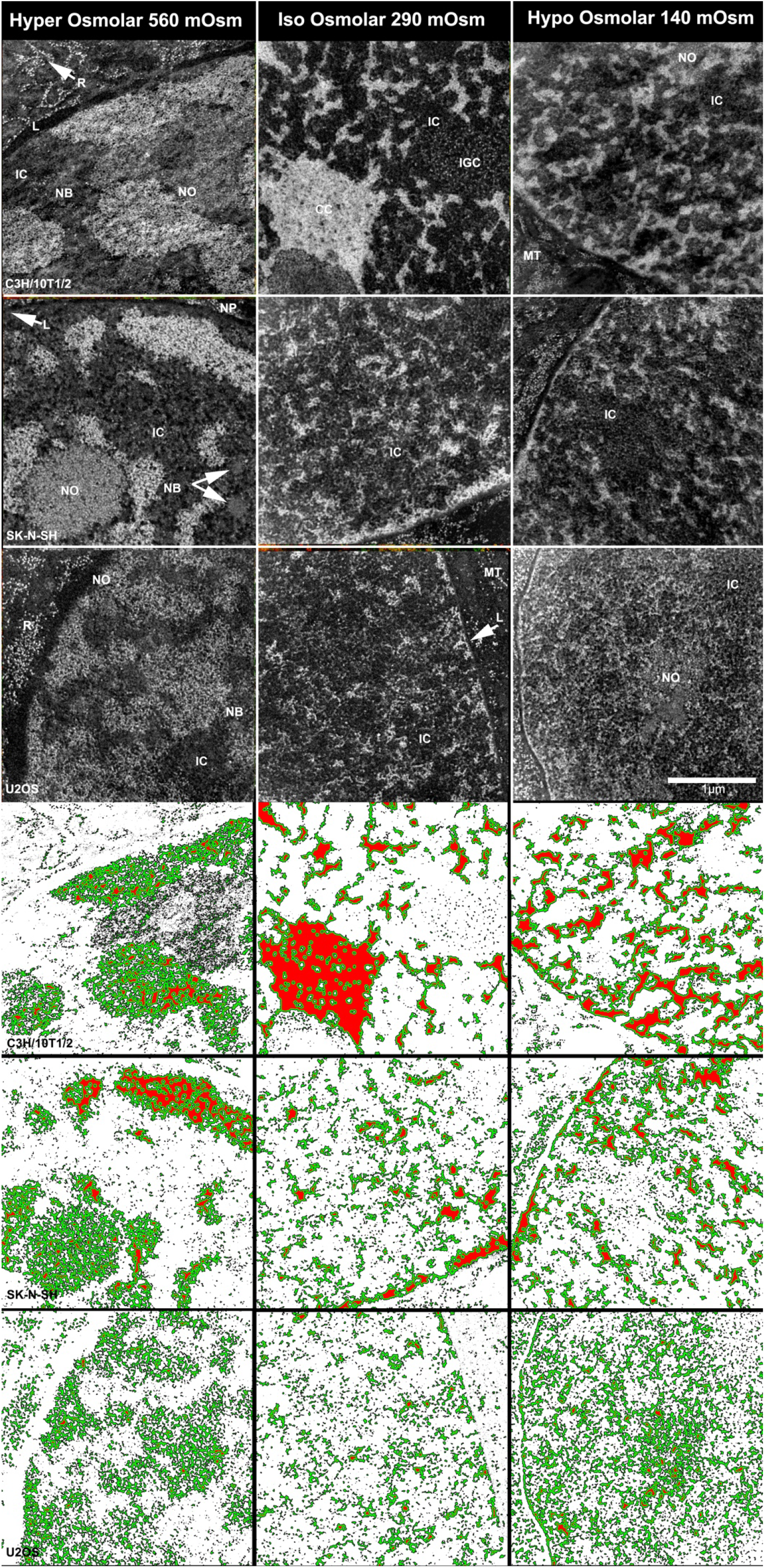
Phosphorous maps (upper 3 rows) and local chromatin thickness maps (lower three rows) of the pictures shown in Figure 3.

**Supporting Online Figure 4:**
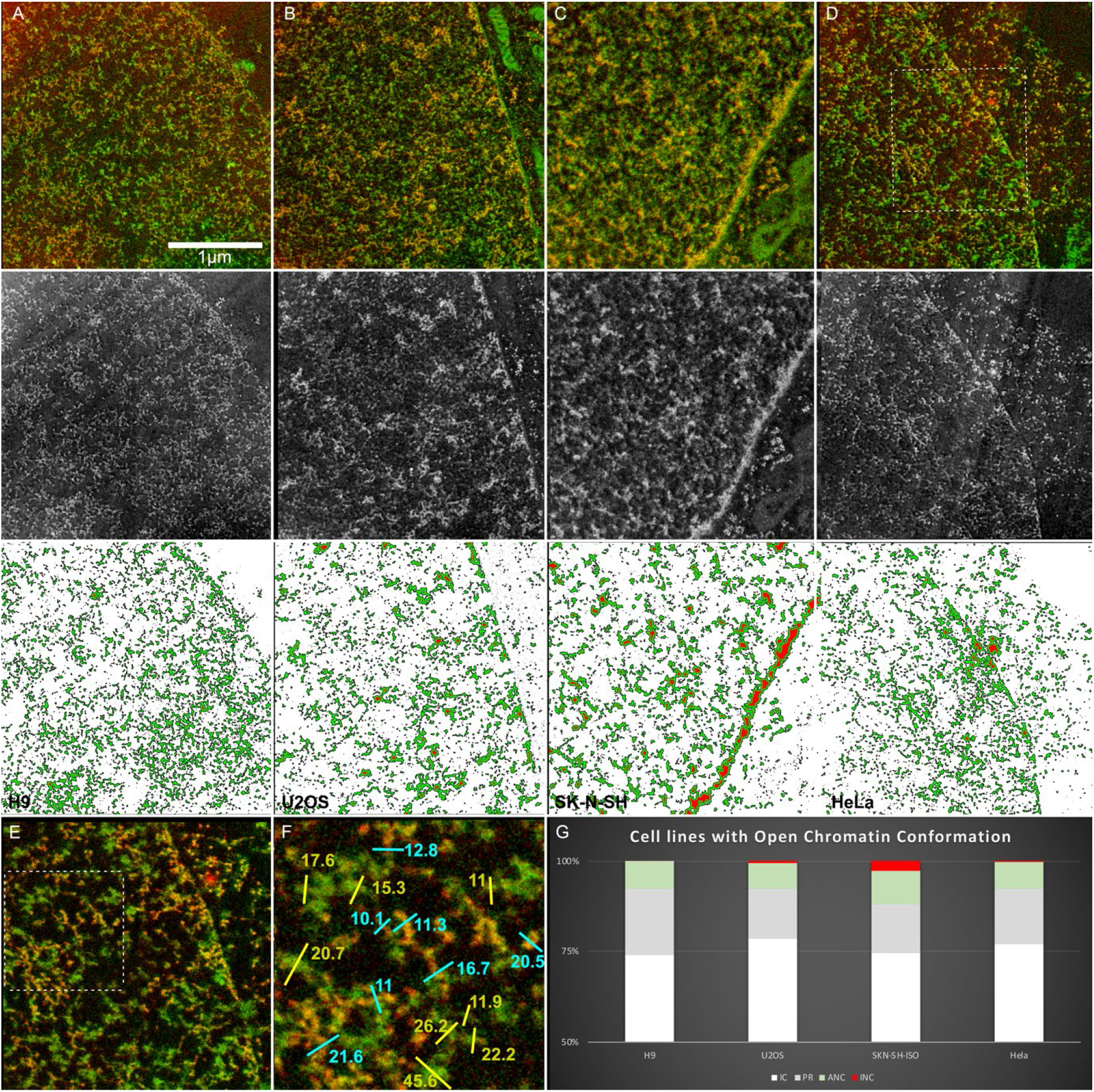
**(A-F):** ESI images (upper), phosphorous maps (middle) and local chromatin thickness maps (lower) of 50nm sections taken from cells with open chromatin conformation. **E-F:** Decondensed chromatin fibers and interchromatin fibrils shown in D at higher magnification and with line scans in F. **G:** Bar charts showing the amounts of Interchromatin Space (white) – chromatin at the interface to the interchromatin space (grey) and proximal (green) and distal (red) chromatin.

**Supporting Online Figure 5:**
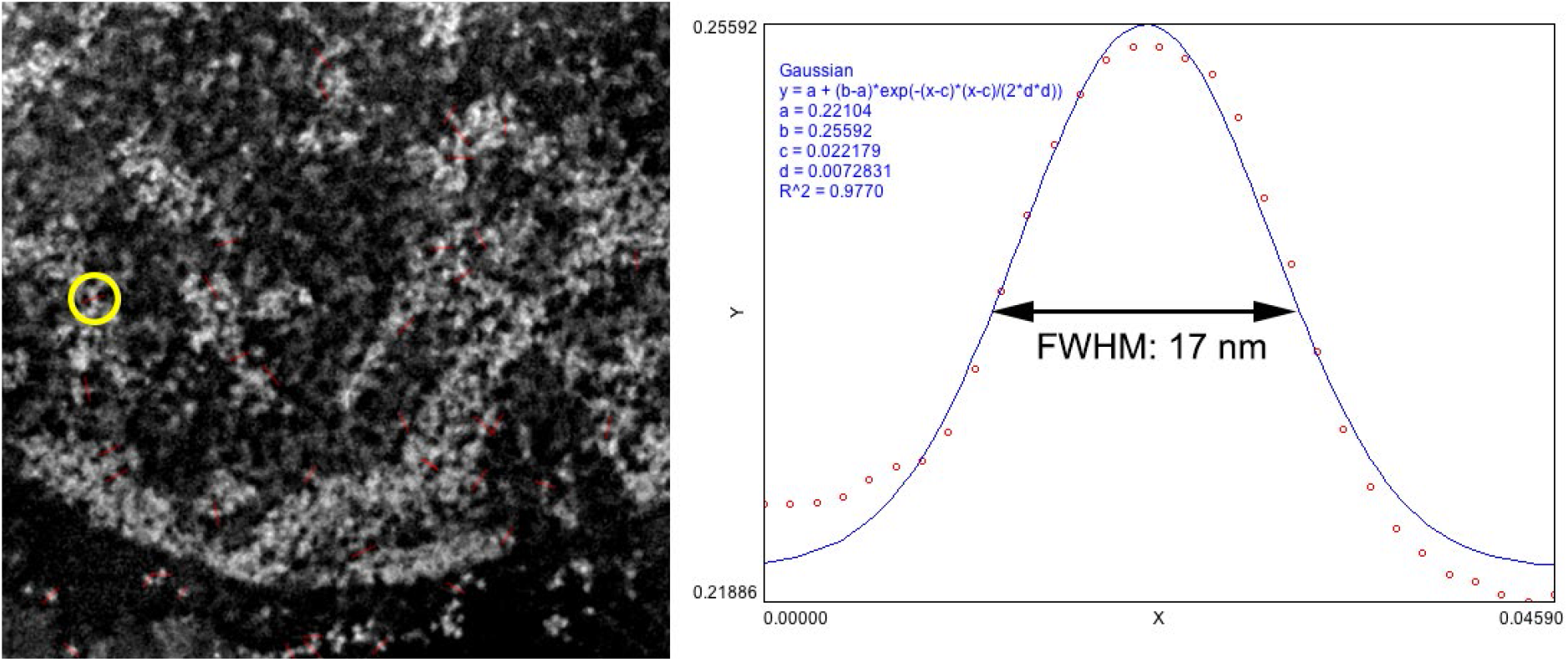
Line scan measurements were performed by drawing a line through a fiber-like structure (left) and determining the intensity profile along this line (right). This intensity profile was then fitted by a gaussian curve. From this fit the full width half maximum FWHM was calculated. FWHMs of ribosomes were used as an internal control in order to compensate for magnification errors that can occur in electron microscopy due to focusing. The mean FWHM of multiple Ribosomes was set to 25 nm and used to normalize the fiber measurements.

**Table S1.**
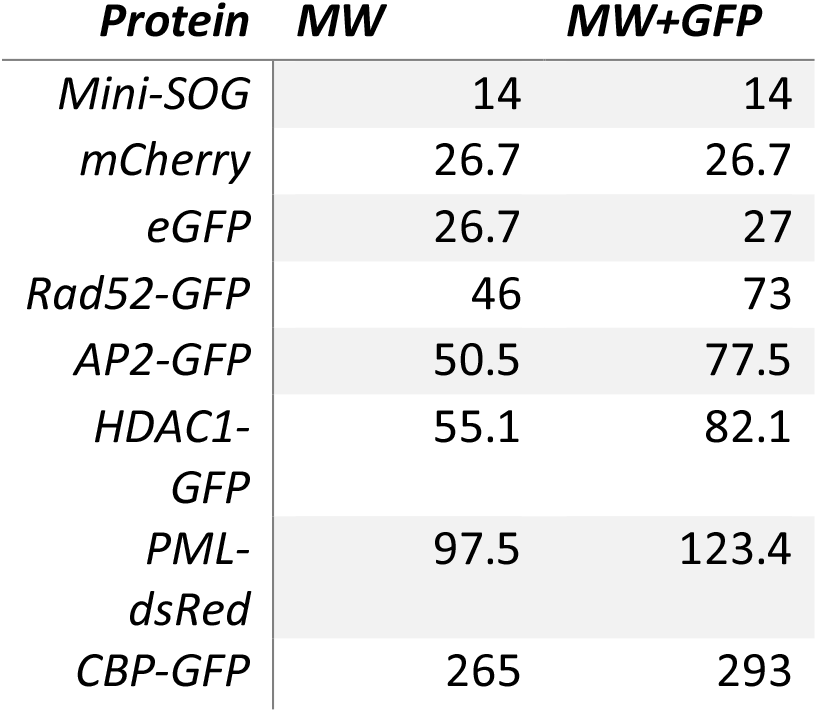

